# The Role of N-acetylcysteine Amide in Acute Graft-versus-host Disease Mouse Model

**DOI:** 10.1101/2025.04.07.647501

**Authors:** Rui He, Wenyi Zheng, Kicky Rozing, Carlos Fernández Moro, Xiaoli Li, Yikai Yin, Weiying Zhou, Samir EL Andaloussi, Svante Norgren, Ying Zhao, Moustapha Hassan

## Abstract

Graft-versus-host disease (GvHD) remains one of the major complications following allogeneic hematopoietic cell transplantation (allo-HCT), resulting in reduced quality of life, morbidity, and mortality in transplanted patients. Clinical strategies to prevent GvHD are frequently associated with off-target effects and dose-related toxicity. Given that oxidative stress is elevated in allo-HCT recipients and contributes to the pathogenesis of GvHD, the current study aims to characterize the role of N-acetylcysteine amide (NACA), a novel antioxidant, as the prophylactic treatment for acute GvHD. Using a murine GvHD model, we found that oral administration of NACA significantly reduced GvHD severity, prolonged survival, and improved the clinical manifestations and integrity of target organs compared to saline or N-acetylcysteine (NAC) treatment. NACA modulated splenic T cells differentiation with an increase in the regulatory (T_reg_) subset and a decrease in the cytotoxic (CD8^+^) subset. Moreover, inflammatory mediators, such as ROS and pro-inflammatory cytokines were downregulated by NACA treatment. In addition, NACA hindered donor T-cell proliferation in the recipients, and restrained Th1 and Th17, but not Th2 polarization. Importantly, NACA did not influence full donor engraftment in bone marrow and spleen. Taken together, our findings provide a new candidate for GvHD prophylactic treatment by targeting oxidative stress that can be easily translated to clinical use.

**Key Points:**

1. NACA provides superior prophylactic effect against aGvHD compared to NAC in an allogeneic transplantation mouse model.
2. NACA treatment neither showed systemic toxicity nor altered the engraftment of the donor cells.

## Introduction

Hematopoietic cell transplantation (HCT) is a therapeutic modality and sometimes is the only treatment strategy for patients with malignant and non-malignant disorders. Despite years of successful HCT, the clinical outcome is still far from satisfactory due to a number of acute and chronic adverse effects including sinusoidal obstructive syndrome, interstitial pneumonia, hemorrhage cystitis, graft-versus-host disease (GvHD), graft rejection and multi-organ failure. GvHD is one of the major complications hampering allogeneic HCT (allo-HCT), causing 10-20% mortality in patients undergoing human leukocyte antigens (HLA)-matched HCT^1^. GvHD is defined as an undesirable immunological response in which the donor T cells encounter host inflammatory antigens and subsequently undergo exaggerated expansion, differentiation, and migration^2^. As a result, the target organs, especially skin, liver and intestines are damaged. Acute GvHD (aGvHD) often occurs within 100 days post transplantation with systemic elevation of Th1, Th17, and sometimes Th2-derived cytokines^3^, and a cumulative incidence of 35% has been reported in a previous study^4^.

Treatment strategies for GvHD are adjusted according to disease severity and presenting symptoms and vary significantly among different clinical protocols. However, most of the clinically used prophylactic regimens rely on calcineurin inhibitors (e.g., cyclosporine and tacrolimus) and anti-proliferative agents (e.g., methotrexate and mycophenolate mofetil)^5^. Recently, post-transplant cyclophosphamide (PTCy) was utilized to prevent GvHD^6,7^ by inducing apoptosis of donor allo-activated T cells and promoting transplant tolerance^8^. Although these strategies can be effective, dose-related side-effects and toxicities may limit their application. For instance, a high level of cyclosporine is associated with graft failure^9^, and PTCy is associated with various cardiac events including left ventricular systolic dysfunction, pericarditis, and arrhythmia^10^. Therefore, new effective prophylactic treatments with superior safety profiles are warranted.

Recently, the role of oxidative stress (OS) in GvHD has gained more attention since it has been recognized as an unavoidable consequence of allo-HCT, which may contribute to T-cell activation and cytokine dysregulation^11,12^. In addition, a dramatic decrease in total and free glutathione (GSH) was reported in mice undergoing allo-HCT compared to those after syngeneic HCT^13^. Prophylactic strategies based on regulating redox hemostasis have shown promising effects in animal models. NecroX-7, a free radical scavenger, was used to regulate aGvHD via mediating the innate immune response. Apocynin, an inhibitor of NADPH oxidase activity, was shown to attenuate GvHD-related mortality by reducing reactive oxygen species (ROS) and levels of proinflammatory cytokines/chemokines^14,15^. Other strategies, focusing on antioxidative enzymes, such as thioredoxin 1^12^ and indoleamine 2,3-dioxygenase 1 (IDO1)^16^, have also been effective in GvHD management.

N-acetylcysteine (NAC) is clinically prescribed as an antidote for paracetamol overdose and as a mucolytic agent and is also known as a classic antioxidant with the ability to scavenge ROS directly and replenish the intracellular GSH pool. NAC has demonstrated anti-inflammatory and immunomodulatory activities in many preclinical settings^17-19^. However, several investigations have recognized that the efficacy of NAC in relieving GvHD severity is highly variable and dose-dependent: a low dose of NAC is immunostimulatory whereas a high dose is immunosuppressive^20,21^. N-acetylcysteine amide (NACA) has a similar chemical structure to NAC, differing only in the substitution of carboxyl for amide, which endows higher lipophilicity, cell-membrane permeability, and bioavailability^22,23^. Moreover, since NACA is converted to NAC after systemic administration^23^, a higher concentration and probably greater immunosuppressive efficacy might be achieved with NACA compared to the same dose of NAC.

As such, we hypothesized that NACA might be an effective prophylactic agent in the context of aGvHD. Our goal was to validate the anti-oxidative and anti-inflammatory potential of NACA as previously shown^24,25^, and to explore its immunosuppressive capability in parallel with NAC.

## Materials and Methods

### Mice

Balb/c (H-2k^d^) and C57BL/6 (H-2k^b^) mice were purchased from Charles River laboratories and maintained in a standard pathogen-free facility. All animal experiments were approved by the Linköping Animal Research Ethical Committee (ID 11257-2020) and were performed according to Swedish Animal Welfare Law.

### Systemic toxicity of N-acetylcysteine amide

Balb/c mice were treated orally with NACA at a dose of 250 mg/kg (bid), for 17 consecutive days. Body weight was monitored, and blood cell counts were measured before and after treatment course using a hematology analyzer (VetScan, HM5). Blood samples were collected. Liver enzymes including alanine transaminase (ALT) and aspartate aminotransferase (AST) levels in serum were detected using commercial ELISA kits (Sigma-Aldrich, MAK052, MAK055).

### Bone marrow transplantation (BMT)

Balb/c mice (12-14 weeks) were conditioned (day −2) by total body irradiation (TBI) at a dose of 800 cGy (divided into two doses, 6-h interval) using a research irradiator (Gammacell 40 Exactor®, MDS Nordion). Mice was intravenously injected (day 0) with 5×10^6^ bone marrow cells (BMCs) and 5×10^6^ splenocytes from C57BL/6 mice for the allogeneic setting, and with same number of cells from Balb/c mice for the syngeneic setting. Clinical GvHD symptoms (weight loss, posture, activity, fur texture and skin integrity) and the survival rate of recipients were monitored. To prevent nonspecific infections, antibiotics (Bactrim, Roche) were added to the drinking water until 14 days post transplantation. Subcutaneous injections of saline were given when severe dehydration and diarrhea were observed (Figure 1A).

**Figure 1.**
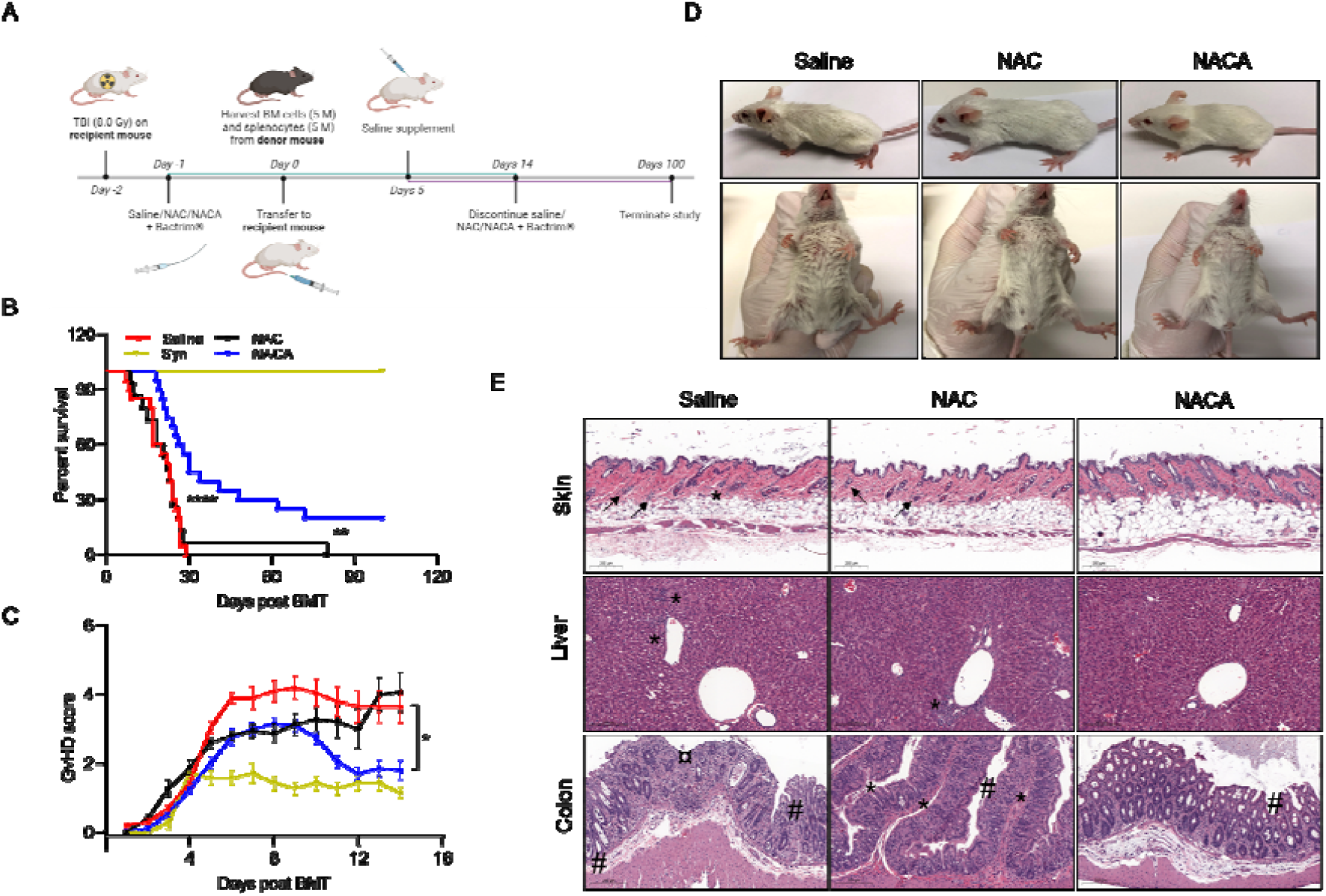
NACA reduces the mortality and morbidity of GvHD. (A) Flow chart of GvHD mouse model establishment and treatment schedule. (B) Survival of Balb/c mice transplanted with 5 × 10^6^ splenocytes plus 5 × 10^6^ BMCs from C57BL/6 mice in saline, NAC and NACA group, or from Balb/c mice in the syngeneic group (Syn). Data was collected from two independent experiments (n = 15). **: p < 0.01; ****: p < 0.0001. (C) GvHD clinical scores were assessed daily post allo-HCT by evaluating the following parameters: body weight, posture, activity, fur texture, and skin integrity were evaluated. Data was collected from two independent experiments (n = 15). *: p < 0.05. (D) Representative images of recipient mice on day 14 post allo-HCT. (E) Representative H&E staining of skin, liver, and colon. Arrow: hair follicle damage or vacuolar changes. *: lymphocytes infiltration. #: apoptosis. ¤: fibrosis with crypts.

### Prophylactic treatments

NACA was provided by Dr Glenn Goldstein (Sentient Life Sciences Inc, NY, USA), and NAC (A7250-25G) was purchased from Sigma-Aldrich. Prior to administration, NACA and NAC were dissolved in saline and neutralized with NaOH, followed by sterilization using a 0.22 μm filter (Millipore, Merck). NACA and NAC were administered orally to recipients post transplantation (day 1-14) at a dose of 250 mg/kg (bid).

### Histopathology

Skin, liver, and colon specimens were obtained from recipient mice seven days after transplantation, fixed in 4% paraformaldehyde (PFA), embedded in paraffin, sectioned, and stained using hematoxylin and eosin (H&E). Slides were scanned using a digital slide scanner (Pannoramic, 3DHISTECH) and images were captured using SlideViewer software (v.2.5.0). The histopathological changes were interpreted by an experienced pathologist.

### Cytokine levels in mouse serum

Serum was collected on day 7 after HCT and the level of interleukin-1 beta (IL-1β) was measured using an IL-1β Mouse ELISA Kit (Invitrogen, BMS6002), and the absorbances were recorded by a microplate reader (Molecular Devices, SpectraMax i3x). The concentrations of 12 other cytokines were detected using the Mouse Th Cytokine Panel (BioLegend, 741043) according to the manufacturer’s protocol. Data were acquired using a MAQSQuant flow cytometer (FACs) and analyzed using LEGENDplex™ Data Analysis Software.

### Determination of reactive oxygen species

Intracellular ROS levels were detected using CellROX™-Green (Invitrogen, C10448C) according to the manufacturer’s instructions. Briefly, single cell suspensions from recipient spleens were permeabilized and labelled with 5 μM CellROX™-Green dye and incubated at 37 °C for 30 min. Cells were then analyzed by a MAQSQuant FACs.

### *In vivo* proliferation assay

Spleens were harvested from C57BL/6 mice and labelled with CellTrace™ Violet (Invitrogen, C34557). Balb/c mice were irradiated at a dose of 800 cGy and transplanted with 8×10^6^ labeled splenocytes. Spleens from recipients were isolated on day 3 post transplantation, data were acquired with FACs (Merck, CellStream™) and analyzed by FlowJo software (v.10.6.2).

### Flow cytometry

Single-cell suspensions were prepared from thymus and spleen and stained for analysis. The conjugated antibodies including Alexa Fluor 647 anti-H-2k^b^ (562832), eFluor 450 anti-H-2k^b^ (48-5958-82), PE anti-H-2k^d^ (553566), PE Cy7 anti-CD4 (100528), FITC anti-CD8 (100706), APC anti-CD25 (102012), PE anti-Foxp3 (126404), eFluor 450 anti-Ki67 (48-5698-82), APC anti-CD62L (17-0621-82), PE anti-CD44 (553134), APC anti-CXCR3 (126512), and PE anti-LPAM-1 (120605) were purchased from BD Bioscience, BioLegend, or eBioscience. Corresponding isotype control staining was carried out in parallel. Surface staining was performed for 30 minutes using pre-mixed antibodies, while intracellular staining was conducted with a Fixation/Permeabilization kit (BioLegend, 426803). Data was collected using a MAQSQuant FACs and analyzed by FlowJo software.

### Intracellular cytokine measurement

Splenocytes from recipients were stimulated *in vitro* with a Cell Activation Cocktail (BioLegend, 423304) at 37°C for 6 h. Intracellular cytokine staining was performed using a Cyto-Fast Fix/Perm Buffer set (BioLegend, 426803). Cells were subsequently incubated with various combinations of the following antibodies from BioLegend or eBioscience: PE-conjugated IL-4 (504104), tumor necrosis factor alpha (TNFα, 506306), IL-17A (12-7177-81); APC-conjugated IL-10 (505010), IL-2 (503810), interferon gamma (IFNγ, 505810). Samples were analyzed using a MAQSQuant FACs.

### Rechallenge of N-acetylcysteine amide

Three long-term (150 days) survivors that showed reactivation of mild GvHD symptoms were treated with NACA at a dose of 250 mg/kg (bid), for one day only. Clinical manifestations were monitored until 17 days after treatment.

### Statistical analysis

Statistics were processed using GraphPad Prism 8.0 software. All data were presented as the mean±standard deviation (SD) unless otherwise stated. The long-rank test was used to compare survival curves, while the 2-tailed nonparametric Mann-Whitney test was applied to the comparison between two groups. Statistical significance was set to p < 0.05.

## Results

### Safety profile of long-term oral administration of N-acetylcysteine amide

NACA has been applied in preclinical settings for more than a decade^26^; however, its safety profile at high doses for continuous administration has not been well studied. Thus, firstly we examined the general toxicity of NACA in Balb/c mice, that would later serve as recipients in our GvHD model. Following oral administration of NACA for 17 successive days at a dose of 250 mg/kg (bid), no significant changes with regard to body weight, blood cell count and liver values were observed (Figure S1), indicating an absence of toxicity. Furthermore, as evidenced in our previous pharmacokinetic study, NACA was rapidly metabolized *in vivo* and eliminated^23^ and thus was unlikely to cause long-term toxicity. Therefore, this dose was considered safe and was used in subsequent investigations.

### N-acetylcysteine amide reduces mortality and morbidity of graft-versus-host disease

The efficacy of NACA in ameliorating aGvHD was evaluated using a classic murine GvHD model with complete MHC-mismatched allo-HCT (C57BL/6 to Balb/c). NACA, NAC or saline was orally administered from the rest day (day −1) after irradiation, until 14 days after transplantation (Figure 1A). Survival profiles were followed for 100 days post BMT. As depicted in Figure 1B, NACA treatment significantly prolonged the survival of mice undergoing allo-HCT with a median survival time of 30 days, while NAC treatment was similar to saline (22 days vs 22.5 days).

GvHD clinical scores, the gold standard for assessing GvHD severity^27^, were recorded throughout the study. The control GvHD mice exhibited a hunched position, decreased movement, ruffled fur, and partially denuded skin at day 14 post allo-HCT, resulting in high clinical GvHD scores. In contrast, a significant decrease in GvHD score was observed in NACA-treated mice, especially from day 12 post transplantation, indicating its protective effect. However, no significant improvement was seen in mice treated with NAC (Figure 1C-D).

Histopathological examination of common target organs in GvHD showed that recipient mice in the NACA group had remarkably reduced severity of tissue damage, with less infiltration of lymphocytes in skin, liver, and colon. In addition, more intact hair follicles and thicker hypodermis layer in the skin, as well as more regular crypt morphology without severe damage in the colon were seen upon NACA treatment (Figure 1E, S2).

### N-acetylcysteine amide promotes engraftment in allogeneic transplanted mice

Engraftment is essential for sustainable and effective hematopoiesis after HCT and is a sign of successful transplantation. It is defined as neutrophil and platelet recovery and is measured by the percentage of donor-derived cells present in the recipients, also known as chimerism^28^. To ascertain the extent of engraftment post transplantation, chimerism assessment and peripheral blood cell counts were performed. We found that donor chimerism in bone morrow on both day 7 and day 14 after transplantation did not vary among groups; however, chimerism in the spleen was slightly enhanced on day 7 following NACA and NAC treatments (Figure S3). Interestingly, the neutrophil count on day 7 was significantly improved in NACA-treated group compared to the control group, whereas no difference was observed regarding the number of platelets, and cell count of other lineages including white blood cells and lymphocytes. Nevertheless, recipients in the NAC group displayed overall lower blood cell counts compared to those in the NACA group, suggesting an inferior statue of hematopoietic reconstitution (Figure S4).

### N-acetylcysteine amide decreases oxidative stress and inflammatory response

Since oxidative damage and inflammation are key factors underlying the pathogenesis of GvHD^29,30^, we next investigated the impact of NACA on ROS accumulation and serum cytokine levels to validate its anti-oxidative and anti-inflammatory activities in aGvHD. ROS accumulation in splenocytes was determined by means of dichlorodi-hydrofluorescein diacetate (DCF-DA) staining at two time points after allo-HCT (Figure 2A-B). Data show that NACA administration significantly reduced ROS accumulation in the spleen compared to that in the control group at both time points, whereas no reduction was observed in the NAC-treated group.

**Figure 2.**
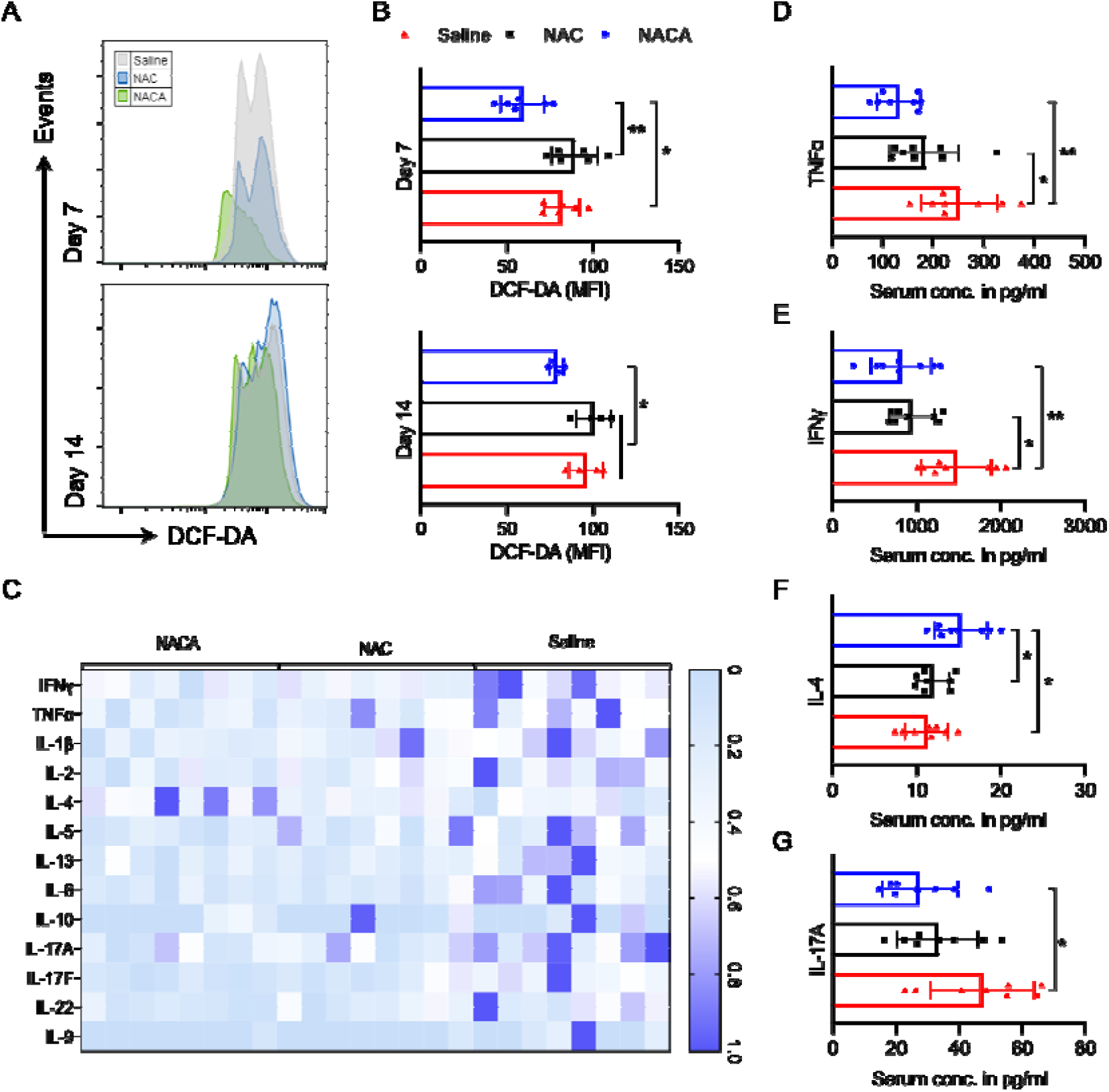
NACA treatment decreases oxidative stress and inflammation in GvHD mice. (A) Splenocytes were harvested on day 7 and 14 after BMT, stained with DCF-DA and analyzed for ROS by flow cytometry. Representative histograms show group differences. (B) Bar charts show individual mean fluorescence intensities (MFI) of each sample with the mean ± SD (n = 6). *: p < 0.05; **: p < 0.01. (C) Heatmap shows serum cytokine levels in individual mouse from each cohort using IL-1β Mouse ELISA Kit and a Mouse Th Cytokine Panel. Samples were obtained from two independent experiments (n=8). (D-G) Statistical analysis of serum levels for TNFα, IFNγ, IL-4 and IL-17A, respectively. Results are presented as individual data with the mean ± SD (n = 8). *: p < 0.05; **: p < 0.01.

Cytokines in peripheral blood after allo-HCT are pivotal mediators and indicators of GvHD^3^, thus serum cytokine levels were quantified. We found that NACA treatment markedly downregulated a number of pro-inflammatory cytokines including IFNγ, TNFα and IL-17A, and increased the level of the anti-inflammatory cytokine, IL-4. However, no effect or a less prominent effect was seen in NAC-treated mice (Figure 2C-G, S5). Together, these results corroborate the superior ability of NACA in downregulating oxidative stress and inflammation compared to NAC.

### Effects of N-acetylcysteine amide on T cell division and proliferation

At the onset of aGvHD pathogenesis, donor T cells traffic rapidly to secondary lymphoid organs after allo-HCT, become activated and proliferate upon exposure to host-derived antigens^31,32^. The spleen is a critical secondary lymphoid organ where donor T cells first react to allogeneic antigens^33^, and is the major object in our study. To assess the immunomodulatory ability of NACA in GvHD, its effect on the expansion of graft-derived T cells was investigated. Briefly, donor splenocytes were labeled with CellTrace™ Violet, inoculated, and analyzed three days post transplantation for violet intensity for cell division. This time point was defined based on a previous report^34^ together with our preliminary results in which donor T cells displayed a moderate division index on day 3 (data not shown). Compared to the control group, fewer divided T cells were observed in the NACA group regardless of the experienced antigen classes, while no changes were found in the NAC group (Figure 3A-B). In combination with our finding about T cell division, the NACA-treated group had significantly lower expression of Ki67, a classic proliferation marker, than the other two groups (Figure 3C-D). Collectively, our data demonstrate that NACA, but not NAC, negatively regulates donor T cell division and proliferation.

**Figure 3.**
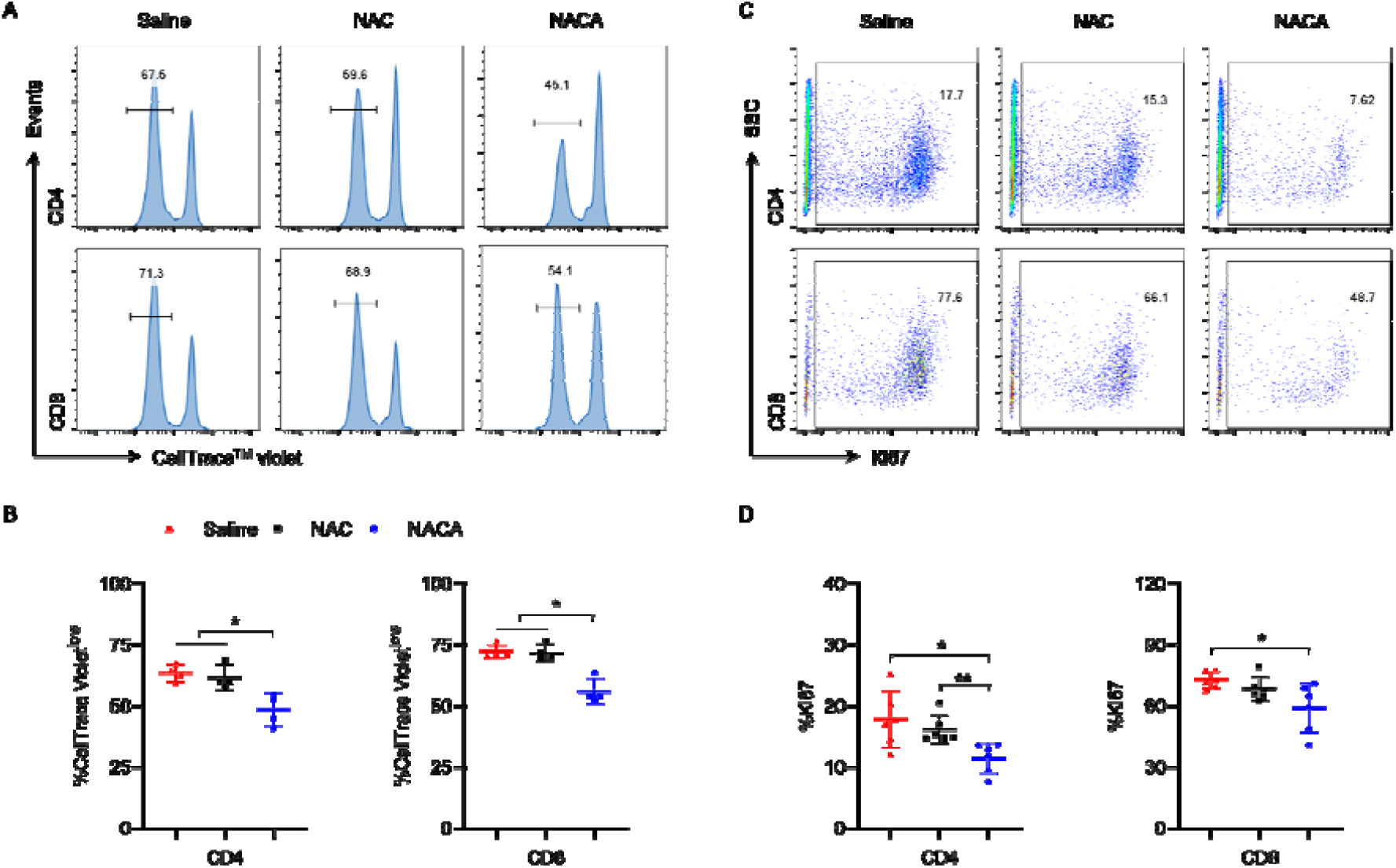
NACA inhibits effector T-cell division and proliferation. Lethally irradiated Balb/c mice were transplanted with 8 × 10^6^ either CellTrace-labeled or unlabeled splenocytes from C57BL/6 mice. All animals were euthanized, and spleens were harvested on day 3. (A) Representative histograms showing donor T cell division. The gated population refers to divided cells with low levels of CellTrace™ Violet (CellTrace Violet^low^). (B) Percentage of divided graft-derived T cells. Results are presented as individual data with the mean value of each group ± SD (n = 4). *: p < 0.05. (C) Representative flow cytometric plots indicate the Ki67 expression in donor T cells. (D) Percentage of Ki67^+^ T cells in each group. Results are presented as individual data with the mean value of each group ± SD (n = 6). *: p < 0.05; **: p < 0.01.

### Impact of N-acetylcysteine amide on T cell phenotype and homeostasis

In addition to T cell expansion, the transition of donor T cells on day 3 was analyzed. Data show that the percentages of effector (CD44^+^CD62L^-^) and central memory (CD44^+^CD62L^+^) CD8^+^ T cells were dramatically increased in the spleen, while the percentages of naïve CD8^+^ T cells (CD44^-^CD62L^+^) were diminished upon NACA treatment in contrast to the control (Figure S6A and C). The same pattern was also observed in CD4^+^ T cells, although the difference in naïve T cells was not significant (Figure S6A-B). Given the linkage of naïve T cells with greater GvHD severity^35,36^, our findings further substantiate the protective role of NACA on aGvHD.

Next, we examined the reconstruction of T-cell subsets in lymphoid organs, an important indicator for GvHD progress. Our results demonstrate that spleen cellularity was replenished to a greater extent in NACA-treated mice compared to those in the control group (data not shown). Furthermore, cytotoxic CD8^+^ cells were significantly reduced in the NACA-treated group, shifting the CD8/CD4 ratio towards a CD4^+^ T cell dominance (Figure 4A-D). In the thymus, NACA significantly increased the CD4^+^CD8^-^ and decreased the CD4^+^CD8^+^ cell populations but had no significant effect on CD4^-^CD8^+^ T cell frequency (Figure 4E-H). Since an increase in CD4^+^CD8^+^ thymocytes is associated with more advanced thymic GvHD^37,38^, the results suggest that NACA might reduce the severity of thymic GvHD.

**Figure 4.**
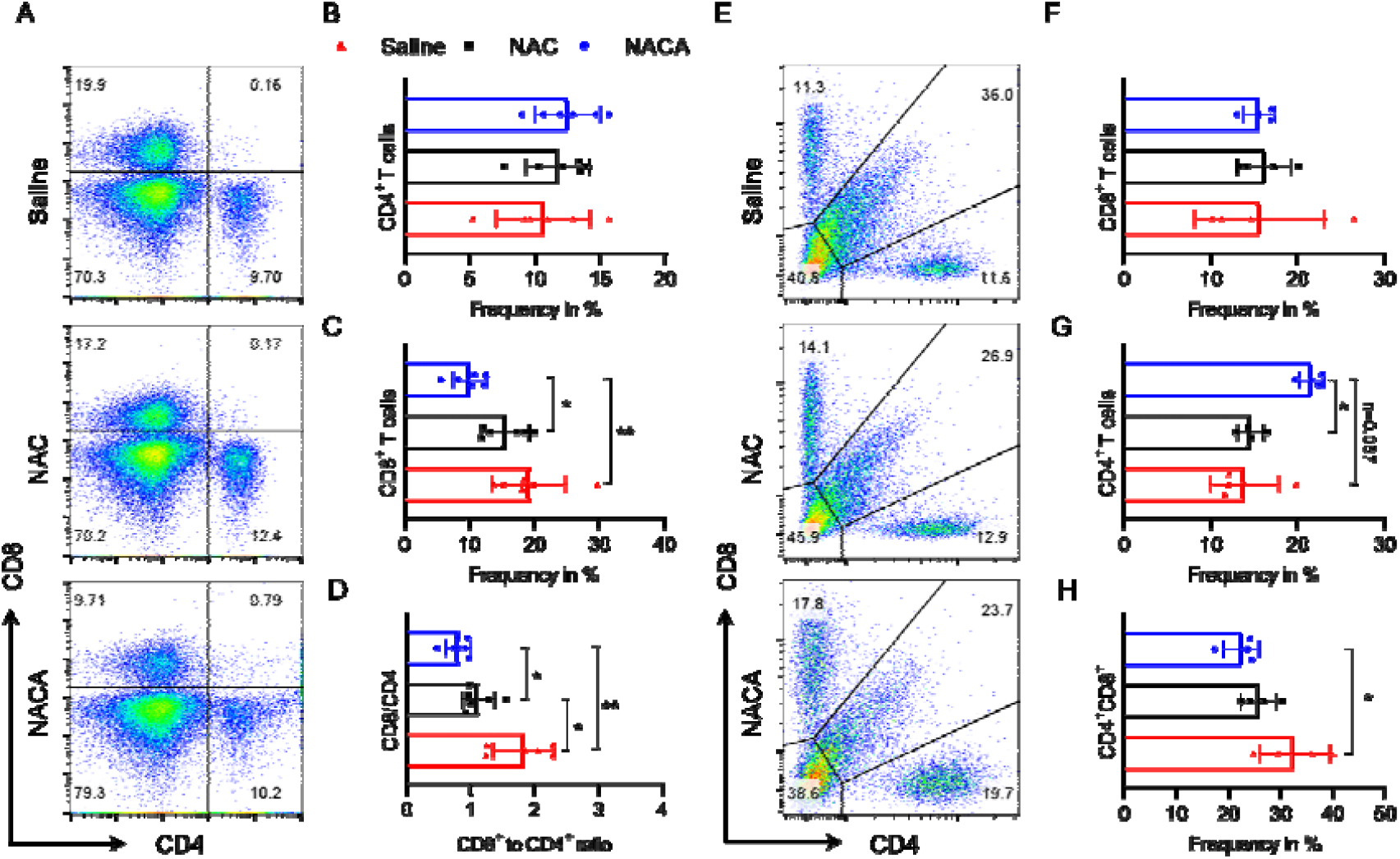
Effect of NACA on T-cell homeostasis. Lethally irradiated Balb/c mice were transplanted with 5 × 10^6^ splenocytes plus 5 × 10^6^ BMCs from C57BL/6 mice. Recipient mice were euthanized on day 7 post HCT, and spleens and thymuses were harvested for analysis. (A) Representative scatter plots show the expression of CD4 and CD8 among gated H-2Kb^+^ splenocytes. (B-D) Quantification of data in A and calculation of the CD8/CD4 ratio. (E) Representative scatter plots show the expression of CD4 and CD8 among gated H-2Kb^+^ thymocytes. (F-H) Quantification of data in E. Results are presented as individual data with the mean value of each group ± SD (n = 4-6). *: p < 0.05; **: p < 0.01.

### Effect of N-acetylcysteine amide on T cell polarization in graft-versus-host disease

GvHD is fundamentally induced by the T-cell allo-response to host antigens^39^. To investigate the effect of NACA on the donor T-cell response to host stimulation, T-cell polarization was investigated in the recipient mice seven days post transplantation. We found that NACA treatment significantly lowered the frequencies of IFNγ and TNFα-producing T cells but increased the proportion of CD25^+^Foxp3^+^ regulatory T cells (Tregs) among donor CD4^+^ lymphocytes in comparison to their counterparts in the other two groups. However, there was no difference between recipients treated with NAC and saline (Figure 5).

**Figure 5.**
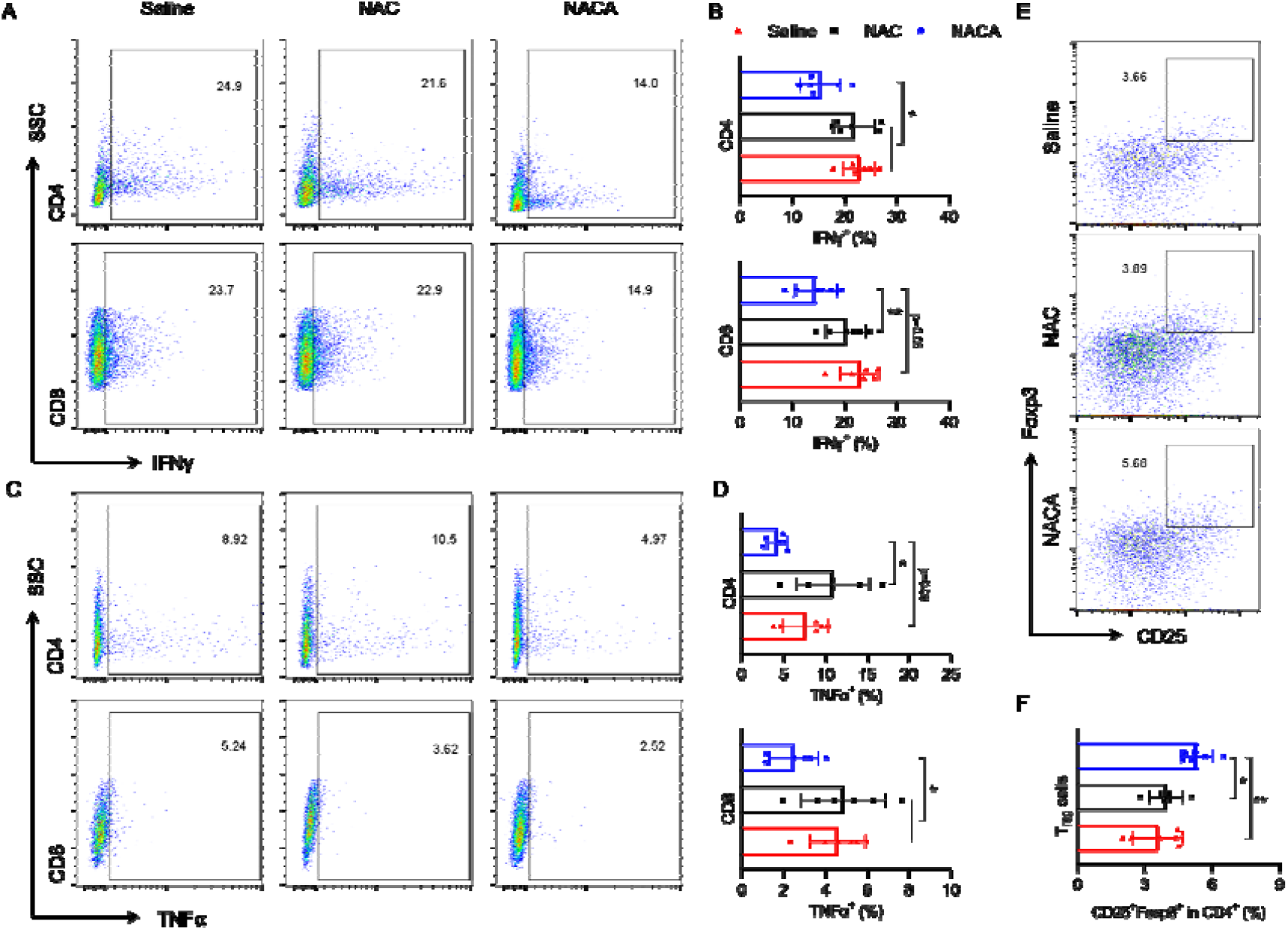
NACA treatment decreases pro-inflammatory mediators but facilities Treg polarization in vivo. (A and C) Splenic T cells from recipient mice were collected on day 7 and underwent in vitro stimulation with phorbol myristate acetate (PMA)/ionomycin for 6 h. The expression levels of IFNγ (A) and TNFα (C) in spleen cells obtained from each mouse were determined by flow cytometry. (B and D) Percentages of IFNγ^+^ (B) and TNFα^+^ (D) T cells among graft-derived splenocytes. (E) Flow plots of representative CD25^+^Foxp3^+^ in donor derived CD4^+^ splenocytes. (F) Frequency of splenic CD4^+^CD25^+^Foxp3^+^. Results are presented as individual data with the mean value of each group ± SD (n = 6). *: p < 0.05; **: p < 0.01.

Meanwhile, we noticed that the proportions of CD4^+^ T cells producing IL-2, although limited to a small positive population of cells, were significantly lower in the spleens of NACA-treated hosts compared to those treated with NAC or saline. Similar results were observed for CD8^+^ T cells (Figure 6A-B). Nevertheless, neither NACA nor NAC altered IL-4 production in either CD4^+^ or CD8^+^ T cells (Figure 6C-D).

**Figure 6.**
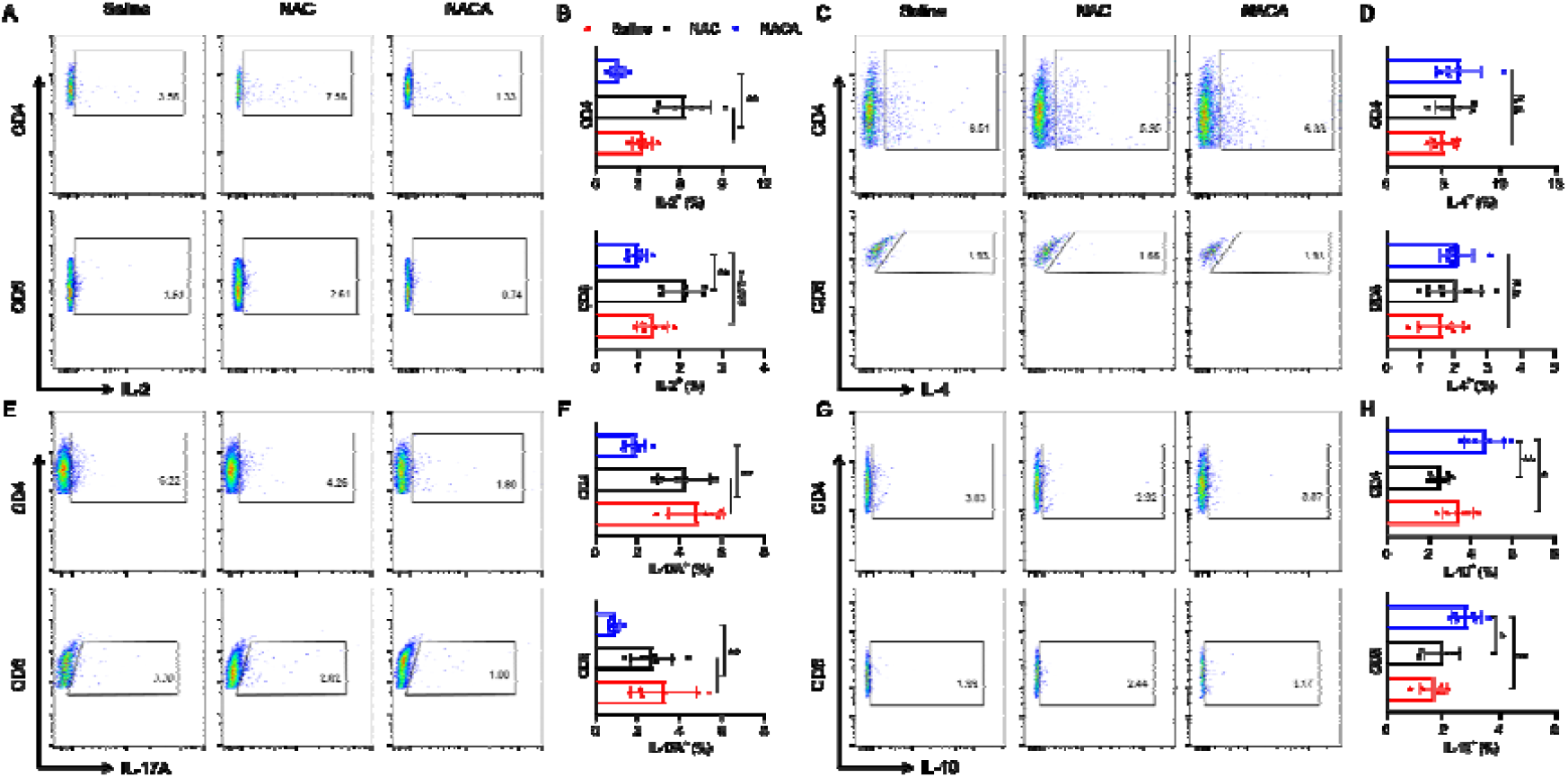
NACA restrains Th1 and Th17 polarization in GvHD. Lethally irradiated Balb/c mice were transplanted with 5 × 10^6^ BMCs and 5 × 10^6^ splenocytes from C57BL/6 mice. Recipient splenic mononuclear cells were isolated for *in vitro* stimulation on day 7 post HCT. Cytokine-expressing T cells were analyzed by flow cytometry after intracellular staining. (A, C, E, G) Flow plots depict one representative mouse in each group regarding IL2^+^ (A), IL-4^+^ (C), IL-17^+^ (E), and IL-10^+^ (G) among gated H-2k^b+^CD4^+^ or CD8^+^ cells. (B, D, F, H) Percentage of determined T cell subsets. Results are presented as individual data with the mean value of each group ± SD (n = 6). *: p < 0.05; **: p < 0.01.

Our results also suggest that NACA could prevent T cells from Th17 differentiation, as IL-17A expression levels decreased dramatically in T cells after NACA treatment, especially in the CD4^+^ subset where the mean percentage was almost halved in contrast to those treated with NAC or saline (Figure 6E-F). Additionally, due to the controversial roles of IL-10 in the development of GvHD^40^, it was of great interest to examine its behavior in the current study. Data show that the fractions of IL-10-producing T cells in recipient mice were significantly higher upon NACA treatment compared to those received NAC or saline as the prophylaxis (Figure 6G-H), implying that IL-10 serves as an anti-inflammatory mediator in our mouse model.

### N-acetylcysteine amide inhibits donor T cell migration to target tissues

Target organ injury in GvHD is driven by the infiltration of effector T cells to the target tissues^41^. CXCR3 is a chemokine receptor that plays a critical role in T cell trafficking, while LPAM-1 (α4β7) is an integrin implicated in the migration of alloreactive T cells to the gastrointestinal tract in GvHD^42^. To investigate the impact of NACA on donor T cell migration, expression levels of CXCR3 and LPAM-1 on donor CD4^+^ or CD8^+^ cells were evaluated.

Results show that mice treated with NACA exhibited a significant decrease in expression of CXCR3 and LPAM-1 in both CD4^+^ and CD8^+^ cells in the spleen compared to their counterparts treated with saline. However, NAC administration did not alter expression levels of the chemokine and integrin in the splenocytes of recipients compared to the control group. Noteworthy, the differences between the NACA and NAC groups were statistically significant (Figure 7).

**Figure 7.**
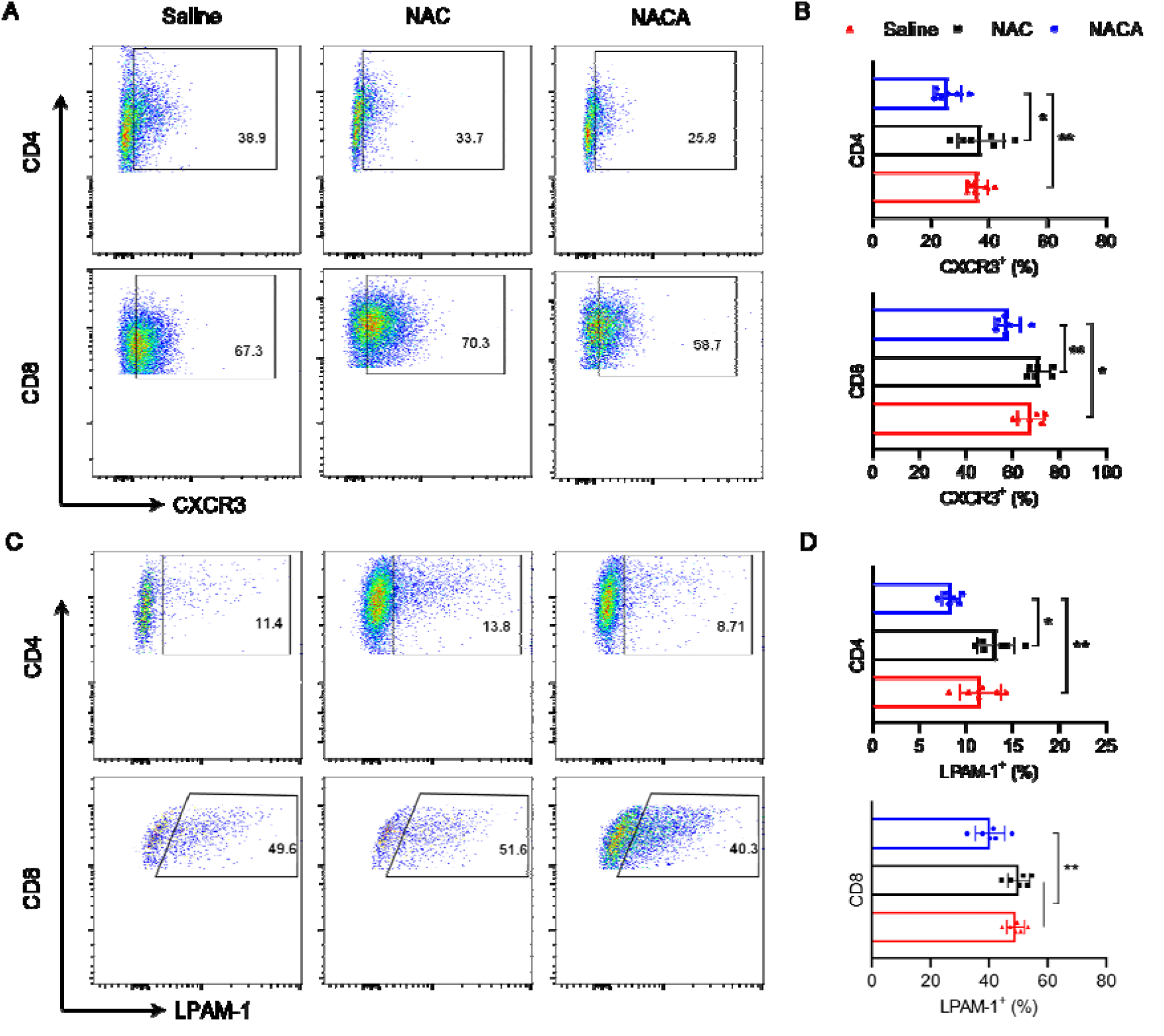
NACA inhibits the migratory ability of donor T cells. Lethally irradiated Balb/c mice were transplanted with 5 × 10^6^ splenocytes plus 5 × 10^6^ BMCs from C57BL/6 mice. Recipient mice were euthanized on day 7 post HCT. (A-B) Recipient splenic mononuclear cells were isolated for analysis of T cell migration surface markers. Representative scatter plots show the expression of CXCR3 or LPAM-1 among gated H-2Kb^+^CD4^+^ or CD8^+^ cells. (C-D) Frequency of CXCR3 or LPAM-1expression in donor CD4^+^ or CD8^+^ cells. Results are presented as individual data with the mean value of each group ± SD (n = 6). *: p < 0.05; **: p < 0.01.

### N-acetylcysteine amide reduced mild GvHD in long-term survivors

Based on the superior ability of NACA to ameliorate the severity of aGvHD, we hypothesized that NACA could be similarly effective in reducing reactivated mild GvHD symptoms in the long-term survivors. Thus, three mice that survived for 150 days after allo-HCT, regardless of previous treatment assignments, received two oral doses of 250 mg/kg NACA with 6-h interval. As depicted in Figure S7A, NACA treatment improved the general condition of these mice, including fur texture, hunched posture, and eye inflammation. Despite there was a slight decrease in body weight two days after treatment, it then remained steady or increased until day 17 post NACA treatment (Figure S7B). In line with the improvement in clinical manifestations, the GvHD score of each mouse declined (Figure S7C). Our results demonstrate the efficacy of NACA in mitigating mild GvHD symptoms seen in long-term allo-HCT survivors.

## Discussion

HCT is a curative treatment for patients with hematological disorders, immune deficiency, or genetic abnormalities^43-46^. However, GvHD remains a significant cause of morbidity and mortality after allo-HCT, warranting efficient strategies for abrogating GvHD to enhance clinical outcome and broaden the use of allo-HCT. The underlying mechanisms are complex and not fully understood; however, the pathogenesis of GvHD is not only associated with systemic inflammation and immunological incompatibility, but is also linked to OS, especially in donor T cells^12,47,48^. In the present study, we propose NACA as a safe and novel antioxidant with the ability to decrease the severity of aGvHD and prolong survival without hampering engraftment.

In line with previous studies^20,21^, we found that NAC had little effect on mitigating aGvHD. Basically, NAC treatment did not improve the survival of GvHD mice, and significantly increased IL-2-producing CD4^+^ and CD8^+^ T cells (Figure 6). IL-2 stimulates the proliferation of cells expressing the IL-2 receptor^49^, and the binding of IL-2 to its receptor may facilitate donor T cell proliferation and expansion^3^, as was observed in our study (Figure 3). Moreover, we found that peripheral lymphocytes were arduous to recover in mice treated with NAC (Figure S4), implying that a worse outcome would occur after transplantation^50^.

In contrast, NACA showed superior ability in alleviating GvHD severity. Firstly, NACA reduced ROS accumulation in the secondary lymphoid organ (Figure 2). Unlike other redox-based prophylactic treatments in aGvHD that usually target certain molecules involved in redox hemostasis^12,14-16,51^, the current study was dedicated to mitigating overall OS, in which multifunctional mechanisms may be involved. One of the most potential pathways is the replenishment of GSH, as our previous report^23^ clearly demonstrated a significant GSH increment after oral administration of NACA. Suh *et al*^13^ reported a rapid depletion of GSH and a notable rise of glutathione disulfide (GSSG) in plasma and liver at an early stage (day 4) post transplantation in GvHD mouse models; however, the substrate cysteine for GSH synthesis was adequate in those mice, implying that NACA, as a cysteine precursor, would be unlikely to replenish GSH and mitigate GvHD. However, NACA is reported to exert its function by reinforcing the activity of glutathione reductase and glutathione peroxidase^52^, that regenerates and make use of the reducing power of GSH, respectively. Besides, NACA can reduce GSSG to GSH via thiol-disulfide exchange without enzymatic catalysis^53^. Therefore, it is believed that NACA acts by replenishing stores of GSH depleted in the early stage of aGvHD via several mechanisms.

In addition, we found that NACA alleviated inflammation in recipient mice by decreasing serum cytokine levels and inhibiting the cytokine producing ability of donor T cells (Figures 2, 5 and 6). A possible pathway through which NACA exerts this anti-inflammatory effect and thus ameliorates GvHD severity may involve regulation of the nuclear factor erythroid 2-related factor 2 - antioxidant responsive element (Nrf2-ARE) axis. Zhou *et al*. reported NACA offers neuroprotection in a traumatic brain injury mouse model via activation of the Nrf2-ARE signaling pathway^54^. In parallel, other studies have identified Nrf2 as an inflammation suppressor through redox control or blockage of proinflammatory cytokine transcription^55,56^. Moreover, studies have proposed the Nrf2-ARE axis as a potent target in pursuit of GvHD treatment^48,51,57^. Such evidence allows us to speculate the rationality of relationship among NACA, Nrf2-ARE and GvHD. However, the exact mechanisms underlying the efficacy of NACA in GvHD need further investigations.

In agreement with our initial hypothesis, significant immunosuppression was observed following NACA treatment. The primary evidence was the marked inhibition of donor T cell division and proliferation (Figure 3). Moreover, NACA impeded the differentiation of donor T cells into Th1 and Th17 subsets but enhanced regulatory polarization (Figures 5 and 6). On day 7 after transplantation, conspicuously higher levels of serum IL-4 were seen in mice treated with NACA (Figure 2), yet the rate of IL-4-producing T cells did not differ from the other two groups (Figure 6). This might be due to a transient increase in serum IL-4 levels since patients receiving tacrolimus and methotrexate for GvHD prophylaxis experienced a similar fluctuation pattern^58^. In addition, we found that donor T-cell trafficking was attenuated based on reduced expression of CXCR3 and LPAM-1 (Figure 7).

As well as inducing GvHD, allogeneic T cells also mediate beneficial effects by eradicating residual leukemia cells, namely graft-versus-leukemia (GvL)^59^. The balance between alleviating GvHD and maintaining GvL is pivotal in the development of new treatments. Although most of the reported strategies regulating redox homeostasis did not interfere with the GvL effect, the fitness of NACA in this regard warrants further validation.

In summary, the present results demonstrate that NACA is safe to use with no sign of systemic toxicity and has a superior prophylactic effect in an aGvHD mouse model without hampering the engraftment of donor cells. The current investigation suggests that NACA can be considered as a promising candidate to be included in the prophylactic treatment of GvHD.

## Acknowledgements

This study was supported by grants from the Swedish Research Council (VR: 2017-00741), the Swedish Childhood Cancer Foundation (Barncancerfonden: PR2017-0083; PR2020-0151), KI funds (2018-02377), and the Karolinska Institutet Center for Medical Innovation (CIMED—20200788) to M.H.. R.H. is supported by the PhD student scholarship from the China Scholarship Council. Dr. Sarah Lang, a medical writer, provided editorial assistance to the authors during preparation of this manuscript.

## Authorship Contributions

R.H., Y.Z., and M.H. conceived the study. R.H., W.Z. K.R., X.L., Y.Y., and Y.Z. performed experiments. R.H. and W.Z. analyzed the majority of acquired data. C.F.M. analyzed the H&E-stained tissue slides. R.H., W.Z. and M.H. wrote the manuscript. S.A. provided the flow cytometer. M.H. supervised the study and acquired funding. All authors participated in the preparation of the manuscript.

## Supporting Information

Figure S1. Systemic toxicity of N-acetylcysteine amide.

Figure S2. Histopathological analysis of GvHD target tissues.

Figure S3. Bone marrow and spleen chimerism after transplantation.

Figure S4. Blood cell counts in GvHD mice.

Figure S5. Cytokine levels in serum.

Figure S6. T cell phenotype in GvHD mice.

Figure S7. Rechallenge with NACA mitigated GvHD symptoms in long-term survivors.

