## supplementary materials for "The Role of N-acetylcysteine Amide in Acute Graft-versus-host Disease Mouse Model"

**Figure S1. Systemic toxicity of N-acetylcysteine amide.** Balb/c mice were administered with NACA at a dose of 250 mg/kg (p.o. bid) for 17successive days. (A) Body weights were monitored, and the weight on day 0 was considered as 100%. (B-C) Mice were sacrificed after the last dose of NACA, and blood samples were collected. After centrifugation, the levels of alanine transaminase (ALT) and aspartate aminotransferase (AST) in serum were detected using commercial ELISA kits (Sigma-Aldrich, MAK052, MAK055). The mice without treatment were taken as controls. (D-F) Blood cell counts were acquired prior to and after treatment course using a hematology analyzer (VetScan, HM5). WBC: whilte blood cells; LYM: lymphocytes; HCT: hematocrit.

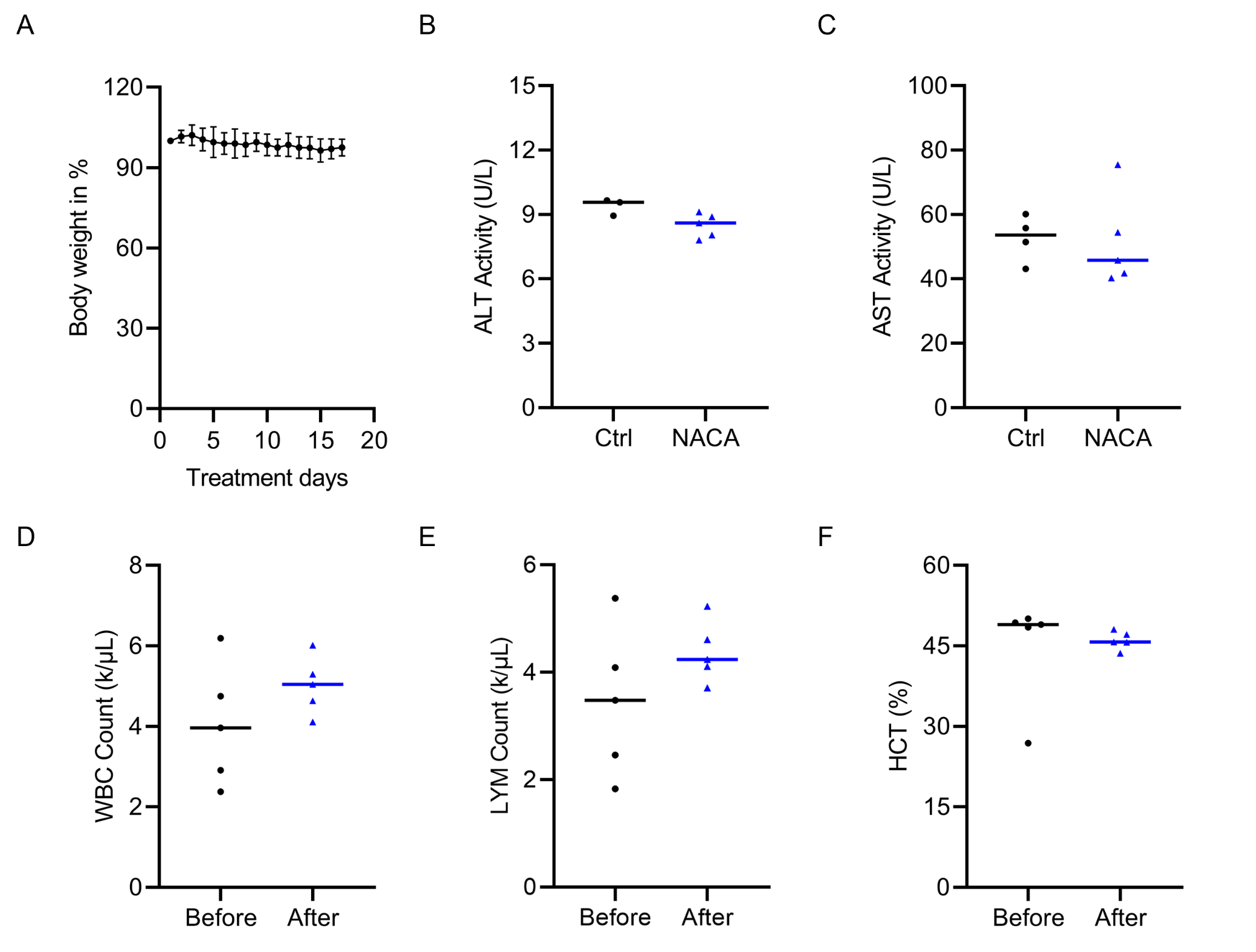

**Figure S2. Histopathological analysis of GvHD target tissues.** H&E-stained tissues were analyzed by an experienced pathologist blindly using QuPath software (version 0.3.2). (A) Representative colon annotations show different classes of mucosal lesion. Dark blue: normal; orange: reactive-fibroinflammatory changes; green: apoptosis; sky blue: fibrosis with crypts. (B) Bar chart shows the percentage of each lesion class. The whole mucosa area is considered 100%. Results are presented as individual data and the mean value of each group (n = 2). (C) Percent of each lesion degree. The whole mucosa area is considered 100%. Results are presented as the mean value of each group (n = 2). Severe: complete fibrosis + necrosis; moderate: fibrosis with crypts; mild: reactive-fibroinflammatory changes + apoptosis; intact: normal. (D) Representative skin annotations show the thickness of each layer. Green: hypodermis; sky blue: dermis; yellow: epidermis. (E) Bar chart shows the thickness of each skin layer. Results are presented as individual data and the mean value of each group (n = 2).

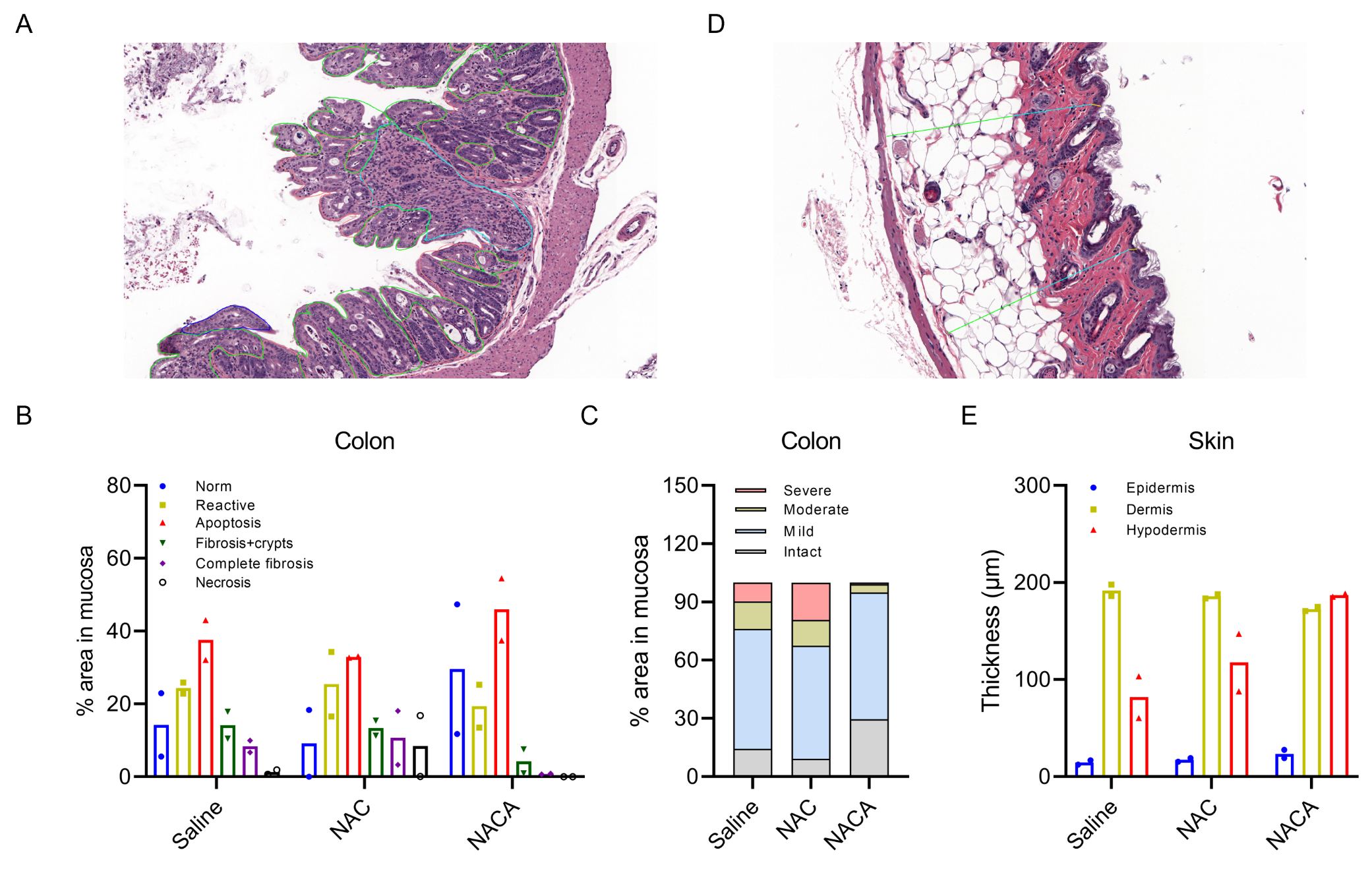

**Figure S3. Bone marrow and spleen chimerism after transplantation.** Balb/c mice received NACA, NAC or saline treatment from day -1 to day 14. After irradiation (day -2), Balb/c mice were transplanted with 5 × 10^6^ splenocytes and 5 × 10^6^ bone marrow cells from C57BL/6 (day 0). On day 7 or day 14 after allo-HCT, recipient mice were sacrificed, and splenocytes and bone marrow cells were harvested. Alexa Fluor 647 anti-H-2k^b^ and PE anti-H-2k^d^ antibodies were utilized for cell surface staining, and data were acquired using a flow cytometer (MAQSQuant) and analyzed by FlowJo software. (A and B) Percentages of H-2k^b+^ cells in bone marrow cells (A) or splenocytes (B) on day 7. Results were presented as individual data with the mean value of each group (n = 10). (C and D) Percentages of H-2k^b+^ cells in bone marrow cells (C) or splenocytes (D) on day 14. Results were presented as individual data with the mean value of each group (n = 5). BM: bone marrow; SP: spleen.

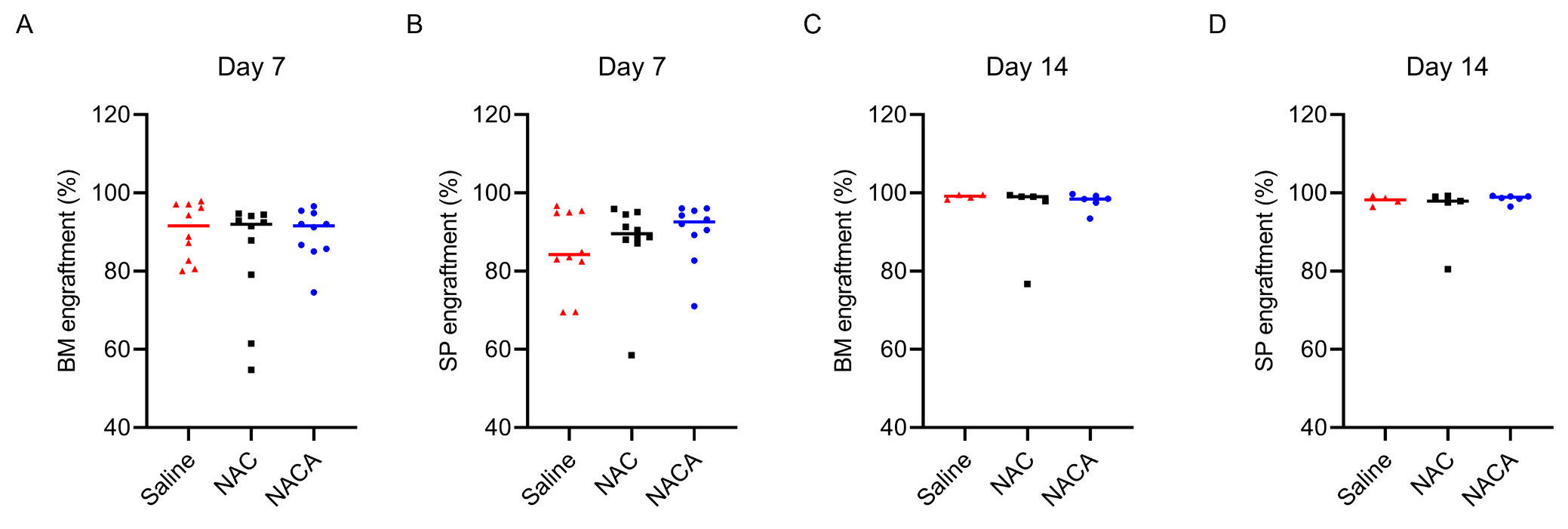

**Figure S4. Blood cell counts in GvHD mice.** Balb/c mice received NACA, NAC or saline treatment from day -1 to day 7. After irradiation (day -2), Balb/c mice were transplanted with 5 × 10^6^ splenocytes and 5 × 10^6^ bone marrow cells from C57BL/6 (day 0). On day 7 after allo-HCT, recipient mice were sacrificed and whole blood was collected by facial vein. Blood cell counts were acquired using a hematology analyzer (VetScan, HM5). WBC: whilte blood cells; LYM: lymphocytes; NEU: neutrophils; PLT: platelet; RBC: red blood cells; HCT: hematocrit. Results were presented as individual data with the mean value of each group ± SD (n = 6). *: p < 0.05; **: p < 0.01.

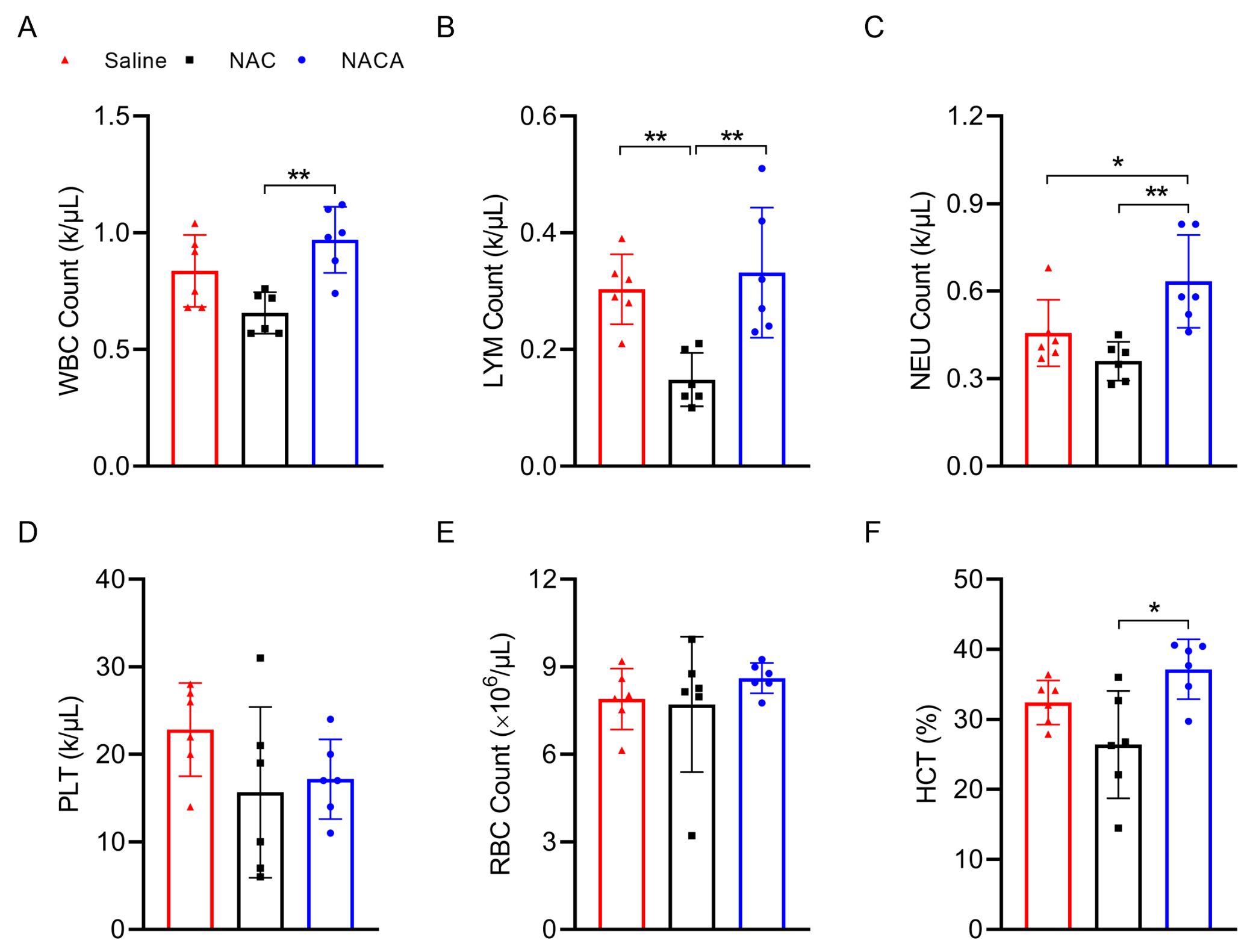

**Figure S5. Cytokine levels in serum.** Balb/c mice received NACA, NAC or saline treatment from day -1 to day 7. After irradiation (day -2), Balb/c mice were transplanted with 5 × 10^6^ splenocytes and 5 × 10^6^ bone marrow cells from C57BL/6 (day 0). On day 7 after allo-HCT, recipient mice were sacrificed and whole blood was collected by heart puncture. After clotting, blood samples were centrifuged at 6000 g at 4 °C for 20 min, and the supernatant (serum) was applied to measurements. (A) The levels of IL-1β were detected by a commercial IL-1 beta Mouse ELISA Kit (Invitrogen), and data were acquired by a microplate reader. (B-I) The cytokine levels were measured using a Mouse Th Cytokine Panel (BioLegend). Data was collected using a flow cytometer (MAQSQuant). Results were presented as individual data with the mean value of each group ± SD (n = 8). *: p < 0.05; **: p < 0.01; ***: p < 0.001.

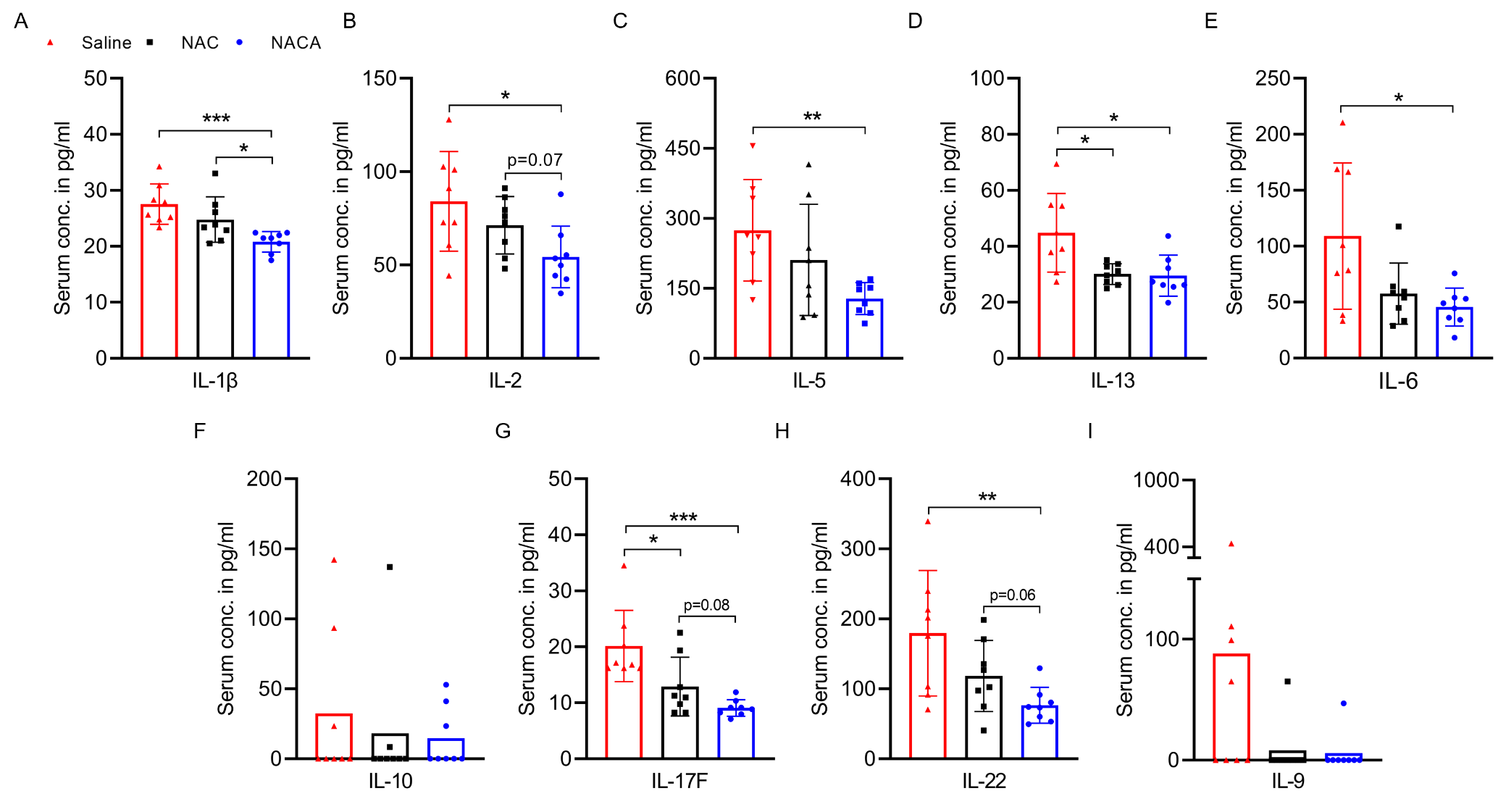

**Figure S6. T cell phenotype in GvHD mice.** Balb/c mice received NACA, NAC or saline treatment from day -1 to day 3. After irradiation (day -2), Balb/c mice were transplanted with 5 × 10^6^ splenocytes and 5 × 10^6^ bone marrow cells from C57BL/6 (day 0). On day 3 after allo-HCT, recipient mice were sacrificed, and spleens were harvested. Single splenocytes suspensions were subjected to a group of surface antibody staining, and data were acquired using a flow cytometer (MAQSQuant) and analyzed by FlowJo software. (A) Representative scatter plots show the expression of CD62 and CD44 in donor-derived T cells. (B and C) Percentages of CD44^+^CD62L^-^, CD44^-^CD62L^+^, and CD44^+^CD62L^+^ in CD4^+^ (B) and CD8^+^ (C) T cells. Results were presented as individual data with the mean value of each group (n = 3). *: p < 0.05.

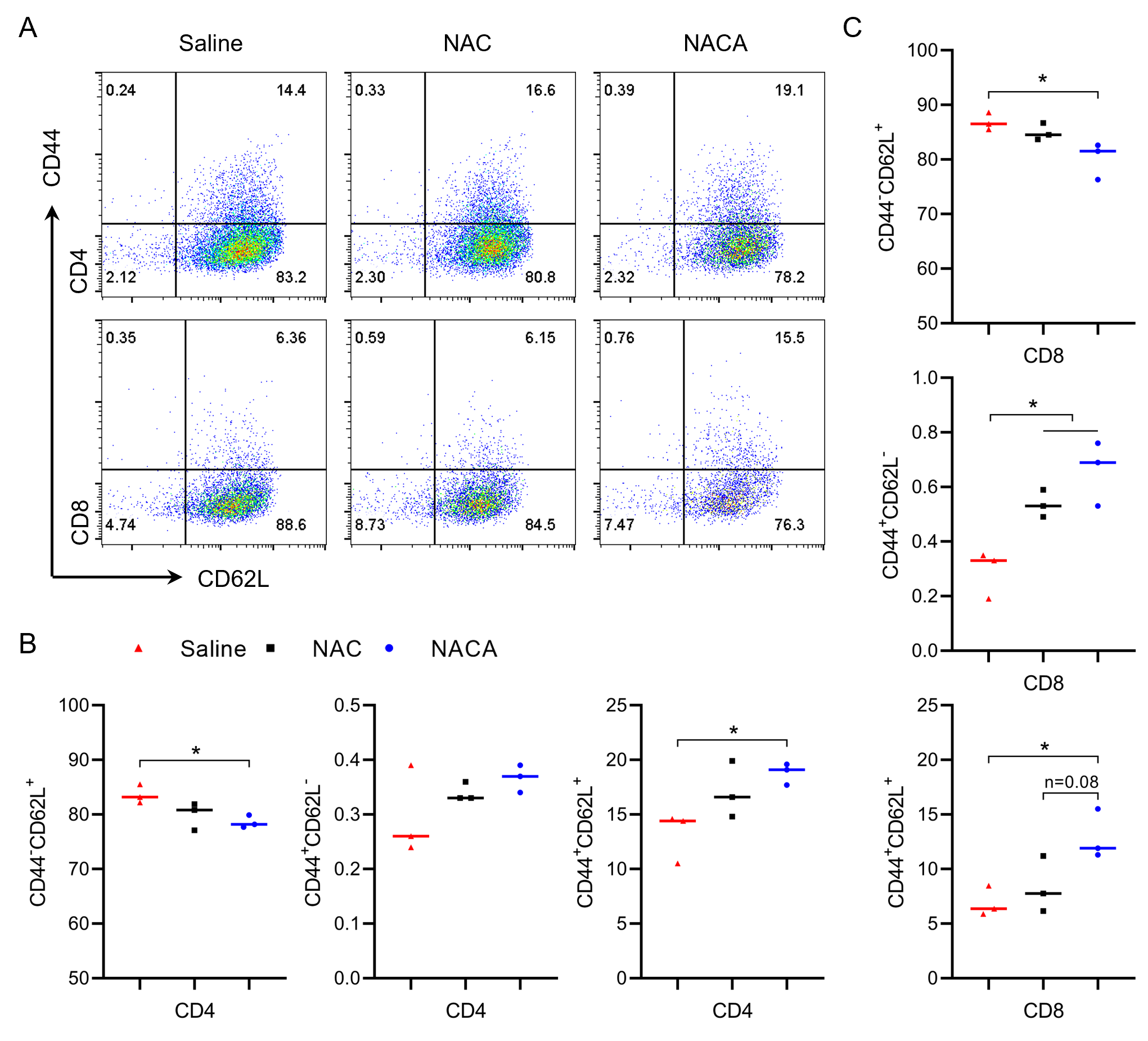

**Figure S7. Rechallenge with NACA mitigates GvHD symptoms in long-term survivors.** Three long-term survivors (150 days after allogeneic transplantation) with mild GvHD symptoms received 500 mg/kg of NACA in two divided doses with a 6-h interval. The mice were followed until 17 days after treatment. (A) Clinical manifestations of each mouse at different time points. (B and C) Body weight (B) and GvHD score (C) of each mouse from three days before to 17 days after NACA administration.

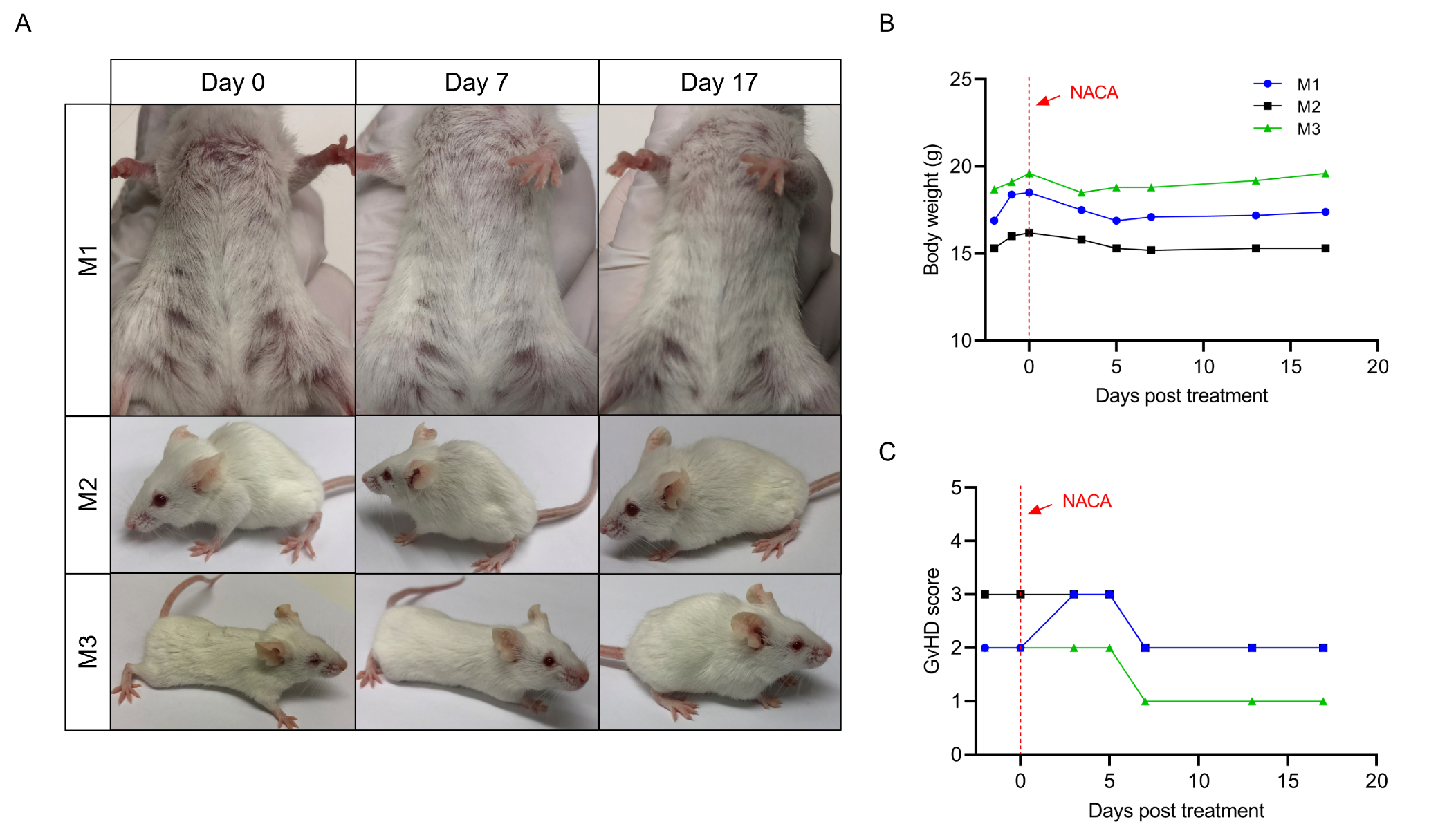
